# Critical length in long read resequencing

**DOI:** 10.1101/621862

**Authors:** De Coster Wouter, Strazisar Mojca, De Rijk Peter

## Abstract

Long read sequencing has a substantial advantage for structural variant discovery and phasing of variants compared to short-read technologies, but the required and optimal read length has not been assessed. In this work, we used simulated long reads and evaluated structural variant discovery and variant phasing using current best practice bioinformatics methods. We determined that optimal discovery of structural variants from human genomes can be obtained with reads of minimally 15 kbp. Haplotyping genes entirely only reaches its optimum from reads of 100 kbp. These findings are important for the design of future long read sequencing projects.

## Background

Long read sequencing using Pacific Biosciences (PacBio) and Oxford Nanopore Technologies (ONT) platforms has profound implications for genomics and genetics [1–4]. In contrast to earlier generations of sequencing technologies, the read length routinely reaches tens to hundreds of kilobases and even up to megabases [5,6].

Long read sequencing leads to more continuous de novo genome assemblies and has advantages for genome resequencing in the context of structural variant (SV) discovery and variant phasing. It enables a more comprehensive detection of genome-wide structural variation, owing to their higher mappability in repetitive regions and their ability to anchor alignments to both sides of a breakpoint [7–9]. SVs are defined as genomic variability of at least 50 bp with a change in copy number or location and include deletions, insertions, inversions, and translocations [10]. It has been shown that 29 000 SVs can be identified per human genome by combining multiple technologies [11], showing that current short read sequencing approaches leave thousands of variants undiscovered. Variant phasing gains from long reads because of the higher chance finding variants inherited from the same haplotype on a single read. Phasing has important implications in determining the pathogenicity of variants and cis-regulation.

To our knowledge, the dependency of SV detection and variant phasing on the read length has not been formally assessed. In this work, we evaluated the influence of the read length on the accuracy and sensitivity of SV detection and on the length of contiguous stretches of phased nucleotides based on simulated PacBio data from a recent assembly of the genome of a Puerto Rican female combining PacBio and Hi-C data using FALCON-Phase [12].

## Results and discussion

Long read resequencing has promising applications for genomics, as it enables direct observation of structural variants and inference of haplotypes [7,9,11]. In this work, we formally assess the impact of increasing read length on the accuracy of structural variant identification and haplotyping of SNVs. While current long read sequencing platforms allow sequencing of tens of kbp to Mbp reads, this comes with a number of disadvantages, requiring more laborious manual DNA extraction from fresh tissue, which may not always be available. Avoiding fragmentation prior to and during library preparation is also essential. Furthermore, striving for ultra long read lengths also seems to reduce the total yield [6]. Due to these limitations and challenges, it is valuable to assess what is the required and sufficient read length to obtain a given accuracy and sensitivity. We approach this problem by simulating PacBio reads based on a recent high-quality phased genome assembly (2.9 Gbp) of the Puerto Rican reference individual HG00733, obtained by combining 75x genome coverage of PacBio long reads with additional long-range information of conformational capture sequencing (Hi-C) (NCBI Assembly identifier GCA_003634875.1). 26 data sets with 40x average coverage and a specific read length starting at 100 bp and up to 500 kbp were simulated in triplicate.

After alignment to the human reference genome GRCh38 with minimap2 [13], we obtained the expected read coverage of ~40x, as assessed with mosdepth [14] (Figure S1). A limitation of our analysis is that we use a fixed read length per dataset, while real long read sequencing experiments produce a long-tailed distribution. We anticipate this simplification is justified to provide approximate guidelines of optimal read length.

The truth set of structural variants was based on a haplotype-specific direct comparison of the HG00733 assembly with the reference genome, resulting in the identification of 25139 variants larger than 50 bp, with a distribution comparable to earlier reports (Figure S2) [7,11,15]. Structural variants from simulated reads were called using Sniffles and compared to the truth set to calculate the precision, recall, and F-measure (Figure 1). An SV is considered concordant (true positive) if it is of the same type and has maximally a pairwise distance of 500 bp between the begin and end coordinates in the test set and the truth set. An additional evaluation filtered variants based on duphold annotation [16], which adds confidence to SVs based on read depth information.

**Figure 1:**
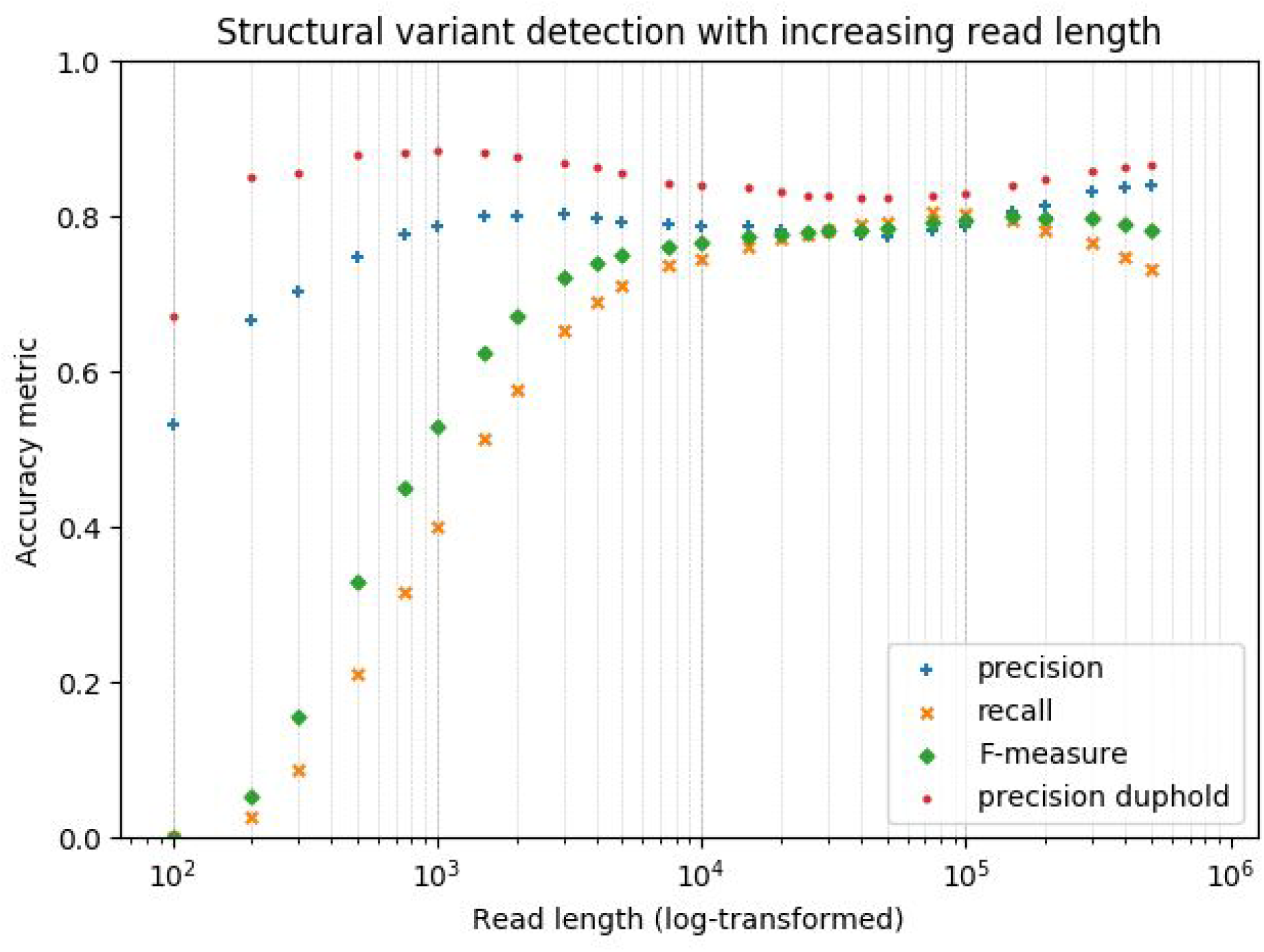
Precision (with and without filtering on duphold annotation), recall and F-measure (y-axis) for structural variant call sets of simulated reads with increasing read length (x-axis). Average of triplicate simulations.

It is worth mentioning that in none of the variant sets from simulated data all variants from the assembly-based truth set are identified, highlighting limitations of the variant caller or suggesting that some variants can only be identified using de novo assembly or read depth based methods. Alternatively, it cannot be fully excluded that the assembly based SV identification contains false positive events, is incomplete, or that coordinates of events that are inferred differently. SV precision reaches its maximum already at reads of 1000 bp, while recall no longer increases substantially after 20 kbp. The F-measure indicates that optimal performance is reached approximately from reads of 15 kbp and longer. Interestingly, recall decreases after 150 kbp (discussed below). Filtering false-positive copy number variants (CNVs) on duphold annotation of read depth changes versus their flanking sequences substantially improved precision, especially for shorter read lengths (Figure 1), while only mildly penalizing recall (Figure S3). As read depth changes are only applicable to CNVs, only the accuracy of deletions and duplications is improved, of which the latter is rarely identified by Sniffles in favor of more common insertions. Replicate simulations showed a high correlation of performance (Pearson’s correlation coefficient >0.97). Notably, as the size of the HG00733 assembly is 89% compared to the human reference genome some genomic content remained unassembled, presumably containing complex repetitive sequences for which longer reads are beneficial for both assembly and SV calling.

Phasing 3.5 million SNVs called from short read sequencing data shows a continuous increase in the length of phase blocks (contiguous haplotyped genomic fragments) with increasing long read length, without reaching a point of saturation within the sizes we tested (Figure 2). Phasing variants across the length of genes is an important application to assess pathogenicity. With reads of 10 kbp about 50% of the genes can be completely phased. This fraction of completely phased genes increases with read length up to a maximum of 90% with reads of 100 kbp or longer, (Supplementary Figure S4). The longest phase blocks are megabases long but are limited by repetitive sequences, regions without identified small variants and structural variation leading to split read alignment. We anticipate that accurate SNV calling methods for long reads would further improve the length of phase blocks, as variants in repetitive sequences cannot be identified by short reads due to ambiguous alignments.

**Figure 2:**
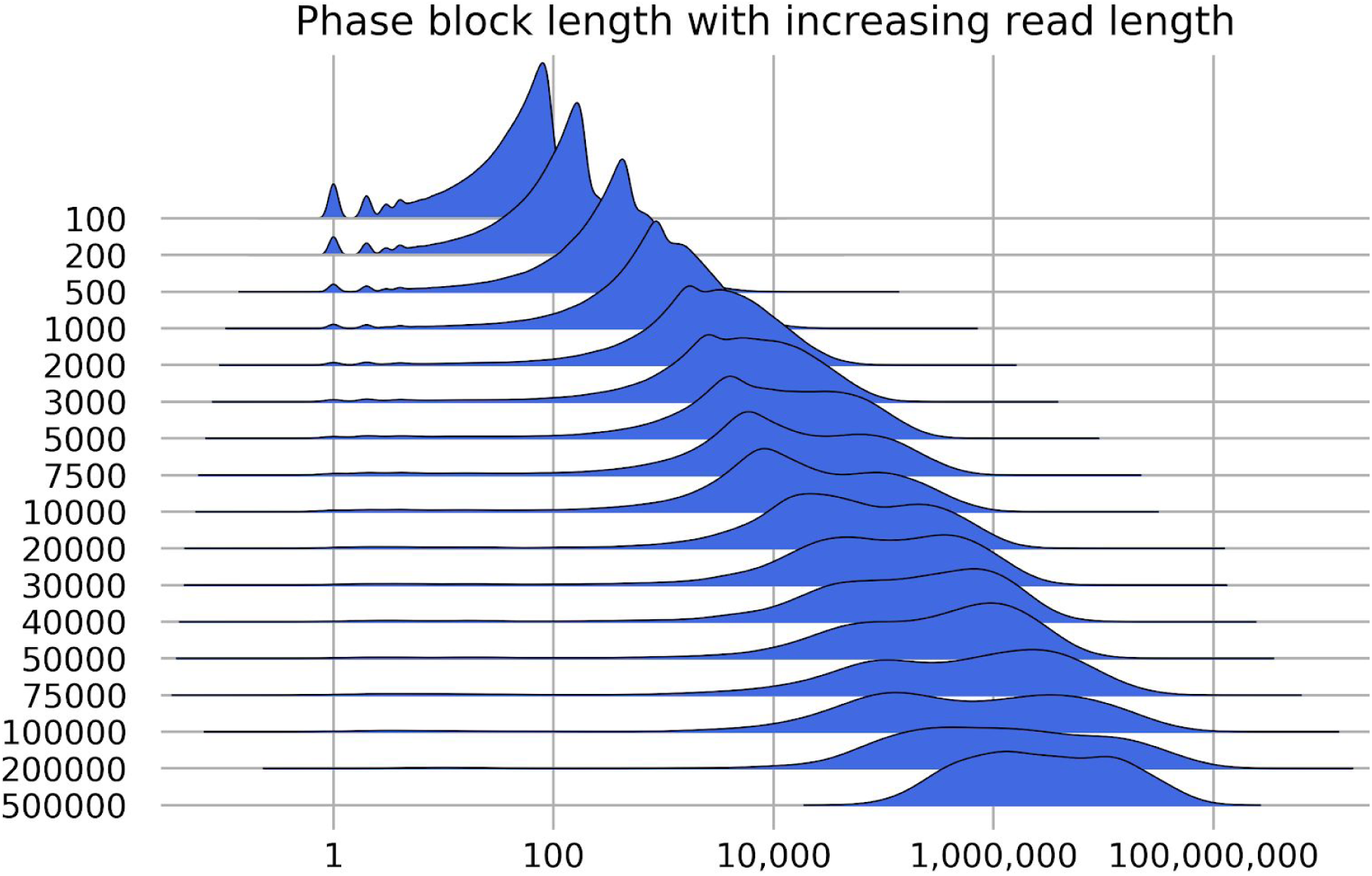
Ridges plot showing the distribution of the length of phase blocks with increasing read length. The x-axis is the genomic size of phase blocks, the y-axis shows the length distribution. Datasets are stacked vertically on separate lines.

With very long reads (> 150 kbp) we surprisingly see that SV recall decreases while precision increases, and to a lesser extent, the proportion of phased genes decreases. As software, including aligners and structural variant callers, was not developed based on such extremely long read sizes, we hypothesize this reduction in performance is an analytic limitation and not due to the increased fragment length itself, e.g. highly complex combinations of SVs with multiple breakpoints per read may be missed, lead to inaccurate alignment or break phase blocks. We can assume this can be mitigated by changing the assumptions of tools used for alignment, variant calling and phasing.

## Conclusions

In the context of human long read resequencing our results show an optimal performance for SV discovery for read lengths of 15 kbp and longer and best phasing across genes from reads of 100 kbp only, crucially guiding the experimental design of future long read sequencing studies.

## Methods

Simlord (v1.0.2) [17] was used for simulation of 40x coverage of PacBio reads with a defined length between 100 bp - 500 kbp (three replicates per length) from a recent assembly of HG00733 (NCBI Assembly identifier GCA_003634875.1, BioProject PRJNA483067) (see commands in supplementary materials). Reads were aligned to the GRCh38 reference genome using minimap2 (v2.14) [13] followed by structural variant calling using Sniffles (v1.0.10) [18]. The obtained read depth was assessed with mosdepth (0.2.3) [14]. The truth set of structural variants was determined using paftools based on the alignment of the HG00733 assembly to the GRCh38 reference alignment using minimap2, after splitting the diploid assembly by haplotype [13,19]. Precision, recall and the F-measure (harmonic mean of precision and recall) are evaluated for SVs with a length larger than or equal to 50th nucleotides using surpyvor [7], which uses SURVIVOR (v1.0.5) for merging VCF files of SVs [20] and cyvcf2 (0.10.0) for parsing VCF files [21]. VCF files with SVs were annotated with the read depth of variants and their flanking sequences using duphold (v0.0.9) [16] and filtered based on the fold-change for the read depth of the copy number variant relative to its flanking regions using bcftools (v1.9) [22]. SNVs from this individual from the 1000 Genomes project [23,24] were phased with the simulated reads by WhatsHap [25], after which contiguous haplotyped segments (phase blocks) were compared to the Ensembl transcript annotation (GRCh38, v95) [26] using bedtools [27]. Data were processed and visualized in Python using pandas (v0.23.4) [28], matplotlib (v3.0.0) [29] and joypy [30]. Commands were parallelized using GNU parallel (v20181022) [31]. Scripts and commands are available at https://github.com/wdecoster/read_length_SV_discovery.

## List of abbreviations

bp: base pair
kbp: kilobase pair
Gbp: gigabase pair
SNV: single nucleotide variant
SV: structural variant
VCF: variant call format

## Declarations

### Ethics approval and consent to participate

Not applicable, as data used by this project is available in the public domain.

### Consent for publication

Not applicable

### Availability of data and material

Results can be replicated starting from the publicly available HG00733 assembly (NCBI Assembly identifier GCA_003634875.1, BioProject PRJNA483067).

### Authors’ contributions

WDC, MS, and PDR designed the study, WDC carried out the informatic analysis. WDC and PDR interpreted the results. WDC wrote the manuscript. All authors read and approved the final manuscript.

## Acknowledgments

The authors wish to thank employees from PacBio and Phase Genomics for making the HG00733 assembly publicly available before publication.

