## Supplementary materials for "Critical length in long read resequencing"

#### Commands

##### simlord

```
simlord -c 20 --without-ns -fl {length} -rr {genome}.fasta --no-sam
simlord-{length}
```

##### Alignment

```
minimap2 --MD -R "@RG\tID:{ID}\tSM:{ID}" -a -t 8 index.mmi {ID}.fastq.gz
```

##### Sniffles

```
sniffles -m {alignment}.bam -v {variants}.vcf -t 4 --genotype
```

##### Variants from assembly to reference alignment (based on [https://github.com/lh3/CHM-eval/tree/master/dip-call](https://github.com/lh3/CHM-eval/tree/master/dip-call))

```
ls hap*.fa.gz|sed s,.fa.gz,,|xargs -i echo minimap2 --paf-no-hit -axasm5 --cs
-r2k -t16 GRCh38.mmi {}.fa.gz 2\> {}.paf.log \> {}.sam.gz | parallel

ls hap*.sam.gz|sed s,.sam.gz,,|xargs -i echo ./sam-flt.js {}.sam.gz \|
samtools sort -m4G -@4 -o {}.bam - | parallel

htsbox pileup -q5 -evcf GRCh38.fa hap*.bam | htsbox bgzip > pair.vcf.gz

./vcf-pair.js -f pair.vcf.gz | htsbox bgzip > final.vcf.gz
```

##### Duphold annotation and filtering

```
duphold --bam {alignment}.bam --fasta {genome}.fna.gz --vcf {variants}.vcf -o
{variants}_dh.vcf

bcftools view -i '(SVTYPE = "DEL" & DHFFC < 0.7) | (SVTYPE = "DUP" & DHFFC >
1.3) | (SVTYPE = "INS")' {variants}.vcf.gz -o {variants}_filtered.vcf
```

##### WhatsHap phasing and phase block statistics

```
whatshap phase --ignore-read-groups SNVs.vcf {alignment}.bam -o
variants_phased.vcf.gz

whatshap stats --chr-lengths chromosome-sizes.txt --gtf {phaseblocks}.gtf
--tsv {phaseblocks}.stats --block-list {phaseblocks}.blocks
variants_phased.vcf.gz
```

### Supplementary figures

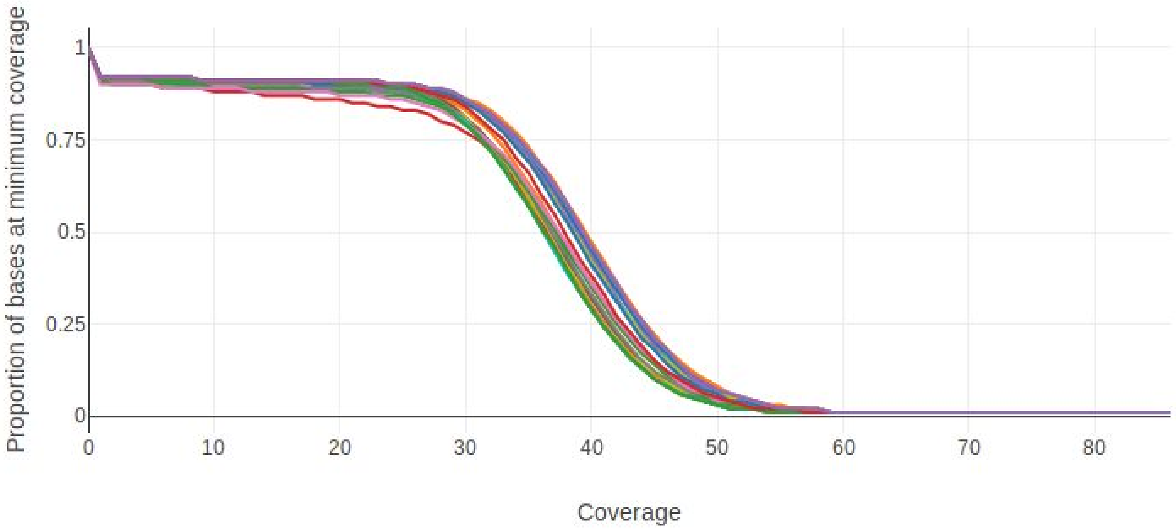


**Figure S1: depth of sequencing coverage assessed using mosdepth**
All simulated read sets show an expected median coverage of 35-40x coverage.

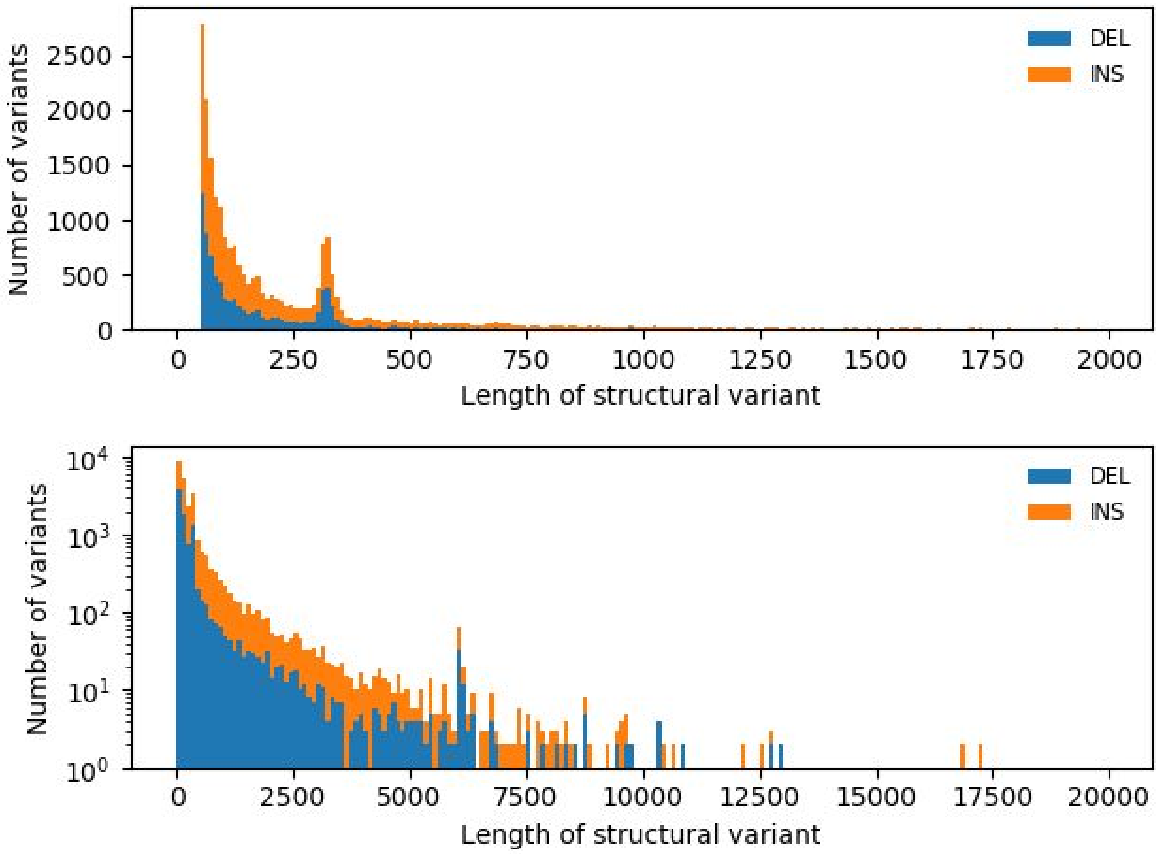


**Figure S2: distribution of the length of structural variants from assembly to reference comparison**

The upper panel shows variants up to 2kb, the lower up to 20kb and uses a log transformation of the number of variants for visualization purposes. Peaks at 300 and 6000 bp correspond to SVs involving respectively Alu and L1 elements.

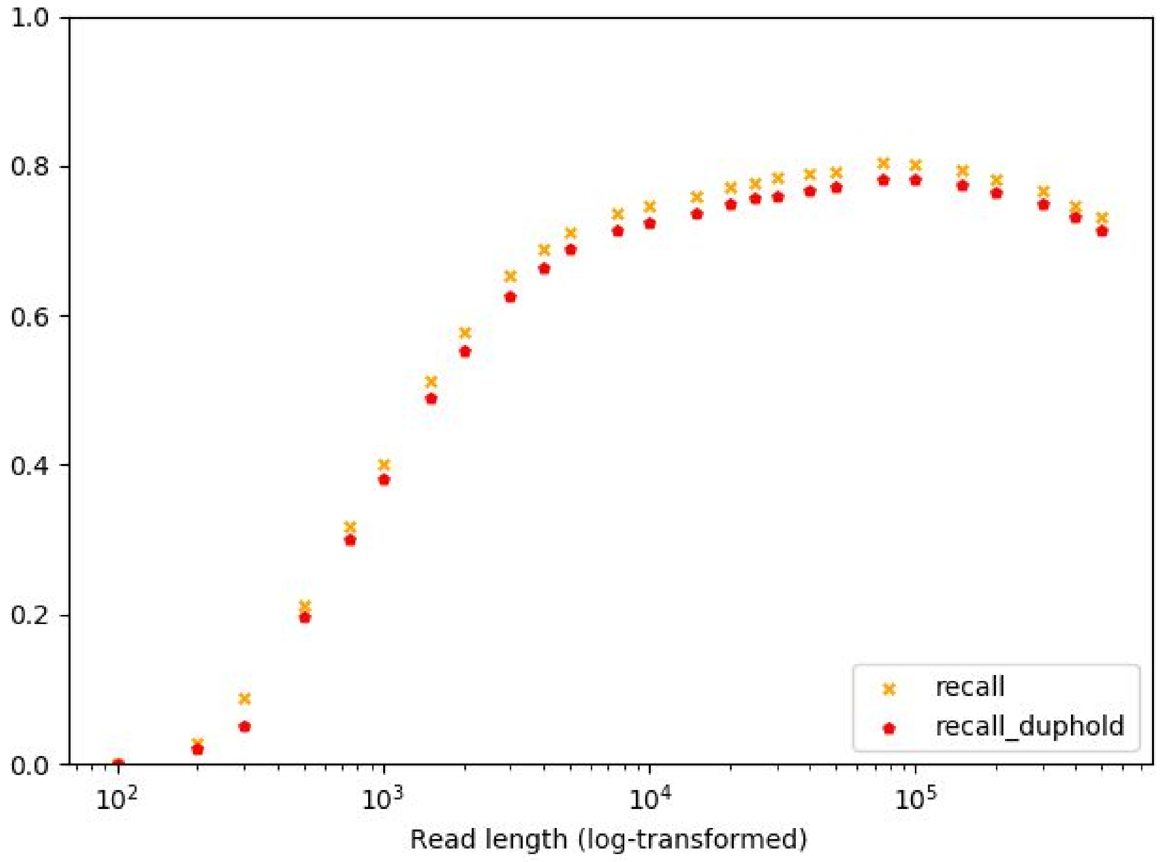


**Figure S3:** A minor loss in recall after filtering on duphold annotation.

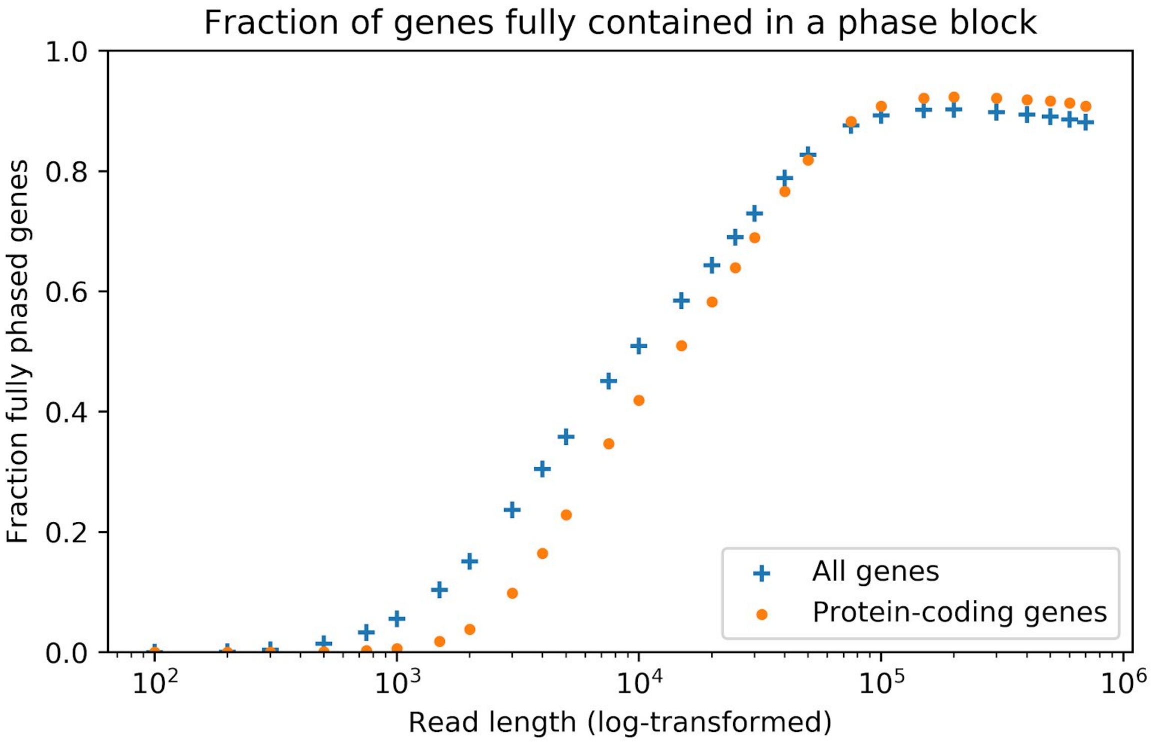


**Figure S4:** the fraction of genes entirely contained in a phase block
